# Detection and Mitigation of Spurious Antisense Reads with RoSA

**DOI:** 10.1101/425900

**Authors:** Kira Mourão, Nicholas J. Schurch, Radoslaw Lucoszek, Kimon Froussios, Katarzyna MacKinnon, Céline Duc, Gordon Simpson, Geoffrey J. Barton

## Abstract

**Motivation:** Antisense transcription is known to have a range of impacts on sense gene expression, including (but not limited to) impeding transcription initiation, disrupting post-transcriptional processes, and enhancing, slowing, or even preventing transcription of the sense gene. Strand-specific RNA-Seq protocols preserve the strand information of the original RNA in the data, and so can be used to identify where antisense transcription may be implicated in regulating gene expression. However, our analysis of 199 strand-specific RNA-Seq experiments reveals that spurious antisense reads are often present in these datasets at levels greater than 1% of sense gene expression levels. Furthermore, these levels can vary substantially even between replicates in the same experiment, potentially disrupting any downstream analysis, if the incorrectly assigned antisense counts dominate the set of genes with high antisense transcription levels. Currently, no tools exist to detect or correct for this spurious antisense signal.

**Results:** Our tool, RoSA (Removal of Spurious Antisense), detects the presence of high levels of spurious antisense read alignments in strand-specific RNA-Seq datasets. It uses incorrectly spliced reads on the antisense strand and/or ERCC spike-ins (if present in the data) to calculate both global and gene-specific antisense correction factors. We demonstrate the utility of our tool to filter out spurious antisense transcript counts in an *Arabidopsis thaliana* RNA-Seq experiment.

**Availability:** RoSA is open source software available under the GPL licence via the Barton Group GitHub page https://github.com/bartongroup.

## 1 Introduction

Antisense RNAs are transcribed from the strand opposite to that of the sense transcript of either protein-coding or non-protein-coding genes. They appear to be widespread in all kingdoms of life and can play distinct roles in regulating gene expression or function. Typically, antisense RNAs are non-coding and expressed at lower levels than sense gene transcripts. However, they can exhibit a range of sizes, and may or may not have 5’ cap or 3’ poly(A) tails depending on whether they arise from either their own promoters, from divergent promoters, or from copying of sense transcripts by RNA-dependent RNA polymerases (see [2] and references therein, [3-5]). In *Arabidopsis thaliana*, for example, the transcription of the Flowering Locus C (FLC) gene is known to be affected by transcription of antisense ncRNAs: COOLAIR [6, 7], a set of ncRNAs antisense to FLC, and COLDAIR [8], antisense to COOLAIR. Both COLDAIR and COOLAIR are associated with different changes in sense strand gene expression at the FLC locus [9]. Antisense transcription is known to affect sense gene expression through multiple mechanisms [2]. During transcription, RNA polymerases may physically interfere with each other if both sense and antisense transcription take place simultaneously. Interference can prevent or slow down transcription (e.g. through RNA polymerase collisions [10, 11]) or force particular isoforms to be produced preferentially [12]. Post-transcriptionally, antisense transcripts can compete with sense transcripts for binding sites [13]. For example, the transcription of the human haemoglobin gene HBA1 is affected when the LUC7L gene on the opposite strand does not terminate, due to a deletion. It produces an antisense transcript that overlaps with HBA1, and which methylates the HBA1 promoter, repressing its expression [14]. In addition, since regions of protein coding genes on opposite DNA strands can overlap, their expression effectively generates transcripts that are, to varying extents, antisense to each other. Such overlapping gene pairs are a common feature of genome organization. We and others have shown that in some eukaryotic genomes tail-tail overlap enables the use of pre-mRNA 3’ processing signals in different registers for genes coded on either strand [15].

Incorporating antisense RNAs into genome annotation and properly quantifying their expression patterns is thus crucial, but challenging. Transcriptome-wide identification of RNAs is currently dominated by RNA-Seq. In this widespread experimental approach RNA is rarely sequenced directly, but instead is fragmented and first copied into cDNA and then copied again, so that libraries of DNA are sequenced. However, the copying of RNA by viral-derived reverse transcriptases is problematic. First, these polymerases exhibit DNA dependent polymerase activity. which can result in copies of the cDNA that can be incorrectly interpreted as antisense transcription. Second, just as reverse transcriptases switch template strand in viral biology, they can similarly switch templates in RNA-Seq library preparation, resulting again, in the interpretation of non-authentic antisense RNAs [16-22]. Historically, in microarray and RT-PCR experiments, this step is known to assign some transcripts to the wrong strand, creating spurious antisense transcripts. Preparing samples with actinomycin D can help to reduce the number of spurious antisense transcripts [18] but can have unwanted side effects [21]. Alternative approaches to make strand-specific RNA-Seq libraries have been developed to mitigate artefacts arising from reverse transcription, however most of these also use reverse transcription [23] and so have similar problems with incorrect assignment. For example, the highly-rated [23, 24] and widely used dUTP protocol for stranded RNA-Seq [25] is known to generate low levels of spurious antisense reads ranging from 0.6-3% of the sense signal [23, 26, 27].Ultimately, the direct sequencing of full-length RNA molecules [28] will overcome many of the problems of distinguishing authentic antisense RNAs. However, currently, reverse-transcriptase based approaches dominate and the extent of spurious antisense RNAs identified in RNA-Seq datasets is rarely exposed.

In this paper, we analyse spurious antisense reads in 199 RNA-Seq experiments, across multiple organisms from both ENCODE [29] and our own work. Our results show that spurious antisense reads are often present in experiments, and can manifest at levels greater than 1% of sense transcript levels. Furthermore, the number of spurious antisense reads can vary substantially between replicates within the same experiment. In some cases, this variation may be sufficient to disrupt downstream analysis of antisense gene expression, by causing spurious antisense counts to dominate the set of genes with high antisense transcription levels.

To detect and correct for wrongly assigned reads, we developed a tool, RoSA (Removal of Spurious Antisense) which calculates an antisense correction factor by identifying subsets of reads where all antisense reads are spurious. We evaluate the effect of using RoSA on *Arabidopsis thaliana* experimental data where varying levels of spurious antisense were present in different replicates. RoSA reduces the overall dependence of antisense counts on sense counts, a key indicator of the presence of spurious antisense. For individual genes with different real and spurious antisense characteristics, RoSA reduces spurious antisense counts while retaining the antisense signal.

## 2 Methods

As noted by Jiang, Schlesinger [26], spurious antisense read counts can be estimated by analysing either ERCC spike-in data, or counts of sense and antisense reads around splice sites. Each approach has different advantages: using spike-ins is simpler and faster, while using spliced reads allows a gene-by-gene estimate to be made. RoSA implements both approaches, in conjunction with pre-processing scripts to generate specialised read counts required by the tool. Once RoSA has an estimate of the levels of spurious antisense, it can adjust the raw antisense counts to account for the incorrectly stranded reads.

### 2.1 RoSA: Removal of Spurious Antisense

Our scripts and analysis code are bundled as a tool, RoSA (Removal of Spurious Antisense), available from the Barton Group’s github pages at https://github.com/bartongroup/km-rosa. RoSA is an R package supported by two python pre-processing scripts, callable from R.

For genes with spliced transcripts which are expressed in the data, RoSA uses the subset of reads from either strand that map across the splice junctions. The antisense reads in this subset are almost certainly spurious, and so RoSA can use the read counts to calculate a gene-specific antisense correction factor (Section 2.2). For genes without spliced transcripts, RoSA uses ERCC spike-in data, if present. Here any antisense read mappings are, by definition, spurious and the ratio of sense to antisense reads mapping to the spike-ins thus provides a global, rather than gene-specific, antisense correction factor (Section 2.3). If ERCC spike-in data is not available, RoSA instead calculates a global estimate of the spurious antisense fraction from the set of spliced reads. Counting all, or spliced-only, antisense reads is not directly supported by existing tools. RoSA’s preprocessing scripts perform these functions. The *make_annotation* script creates an antisense annotation (as gtf) from a standard annotation (as gff or gtf), which can then be used to generate antisense read counts via a standard read counting tool (Section 2.4.1). RoSA adjusts these raw counts to produce corrected antisense counts (Section 2.4). The *count_spliced* script generates sense and antisense counts of reads at splice junctions, used when estimating spurious antisense from spliced reads. The script takes a standard annotation (as gtf/gff) and corresponding alignment (as bam) and outputs counts of spliced sense and antisense reads to a designated output file.

The RoSA R code takes as input datasets containing several different read counts, for each replicate:

1. Full read counts by gene
2. Antisense counts by gene (via the *make_annotation* script)
3. At least one of:

a. Spike-in sense and antisense counts
b. Spliced sense and antisense counts (via RoSA’s *count_spliced* script)

RoSA calculates and returns antisense:sense ratios for the spike-in data, or spliced read data, or both, and, for each gene and replicate, outputs new read counts values corrected for spurious antisense. RoSA also plots antisense versus sense counts of the original and corrected data, by replicate.

### 2.2 Using spliced reads

RoSA’s main approach to estimating spurious antisense is to use spliced reads within the main dataset. Reads which map antisense to a multi-exon gene, and that also show the same splicing pattern as spliced sense-mapping reads are almost certainly spurious, as the splicing motif (canonically GU-AG) will be incorrect on the opposite strand (see Figure 1 for an example). An estimate of spurious antisense can be calculated by considering only spliced reads whose splices match annotated splice sites (*splice-matched reads*), and, as with the spike-ins, calculating the ratio of antisense to sense reads.

**Figure 1:**
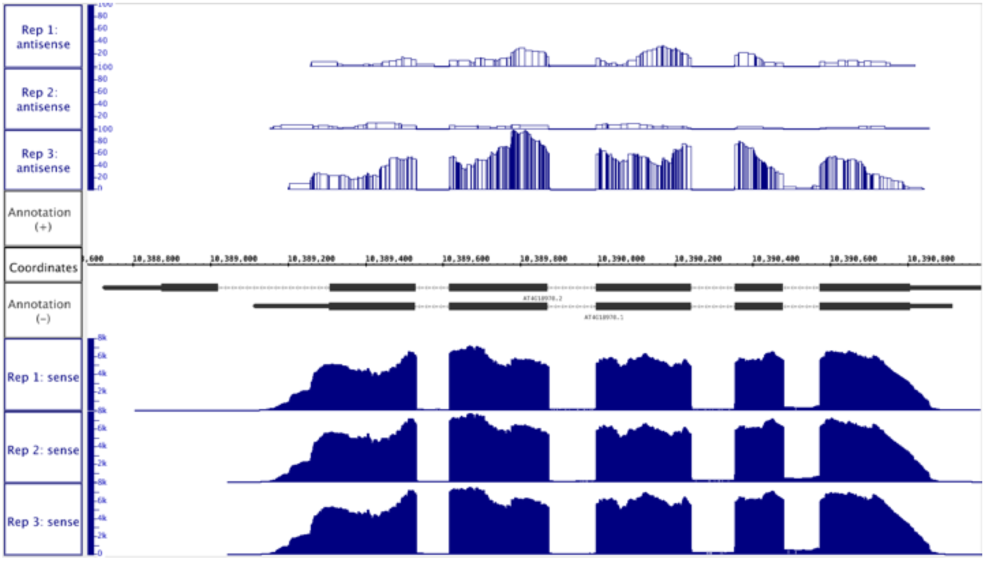
An example of spurious antisense reads displaying the same splicing structure as the sense strand. The reverse strand gene AT4G18970 is strongly expressed in all 3 replicates (bottom tracks). Spurious antisense can also be seen in all replicates (top tracks), with splice points in the antisense signal matching splice points in the sense signal. Furthermore, the level of spurious antisense varies noticeably between replicates. (Figure generated by IGB [1])

Splice-matched reads are identified by first filtering all spliced reads in the data. In a bam file of aligned reads, spliced reads have a CIGAR string containing ‘N’, indicating a skipped region. SAM processing tools such as sambamba [30] support filtering on the CIGAR string and can extract spliced reads rapidly. A second filtering step pulls out only those reads whose splice locations match at least one intron in the annotation, by processing each read in turn, identifying the spliced positions (based on the read location and the CIGAR string) and checking the annotation for a matching intron.

Finally, the strand of each spliced read can be determined from its flag field value [31], and compared to the strand of the matching intron(s). Reads on the same strand as the intron(s) are counted as sense reads, and remaining reads as antisense reads. Since spurious antisense reads are misallocated sense reads, the number of antisense splice-matched reads assigned to a gene is strongly positively correlated with the number of its sense splice-matched reads (see Section 3). The ratio of antisense:sense counts on the splice-matched reads thus gives a simple global estimate of the level of spurious antisense across the whole dataset. Using spliced reads has the advantage that an antisense:sense ratio can be calculated on a gene-by-gene basis, for any spliced gene. Genes without any spliced reads fall back on the global estimate, calculated either from the spike-ins (see Section 2.3) or the spliced reads.

### 2.3 Using ERCC spike-ins

An alternative approach to estimating spurious antisense is to use ERCC spike-in data. Developed by the External RNA Control Consortium (ERCC) [32], the ERCC spike-in controls are synthetic RNA transcripts that are added to RNA-Seq experiments to act as controls [33]. The 92 spike-ins are designed to mimic a range of eukaryotic mRNA characteristics, varying in length, GC-content and concentration, with a 20bp poly-A tail. They have minimal sequence similarity with known eukaryotic transcripts. Since the spike-ins are synthetic, they are unidirectional, and so any reads assigned as antisense to a spike-in can be assumed to be spurious. As the spike-ins are present at a wide range of concentrations, they are detected with a wide range of read counts, permitting an estimate of the ratio of antisense to sense read counts on the spike-ins to be calculated, which can then be used to estimate the contribution of spurious antisense transcripts across the full dataset. Obtaining sense and antisense counts for the spike-ins is straightforward. First we align the reads to the spike-ins (using the spike-in annotation file ERCC92.gtf, available at https://www.thermofisher.com/order/catalog/product/4456739) and then count reads, using a strand-aware read counting tool such as featureCounts [34], HT-SeqCount [35], etc. Now averaging the spurious antisense:sense ratio across all of the spike-ins gives a global estimate of the spurious antisense, in just the same way as for the spliced reads.

### 2.4 Mitigating spurious antisense

Having identified high or differing levels of spurious antisense in an RNA-Seq experiment, we also want to correct for the incorrectly assigned reads so that true differential expression calling can be performed. The ratio of spurious antisense:sense read counts can be used as a simple correction factor. Defining *r* as the ratio of spurious antisense:sense and *S* and *A* respectively as the number of sense and antisense counts for a gene, the number of spurious antisense read counts *AS* is estimated for each gene as:

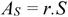

Then the antisense count can be corrected to account for the spurious antisense by taking *A* - *A_S_*. This correction simply adjusts read counts for each gene, and does not identify specific reads as incorrectly assigned, so pile-ups cannot be adjusted. Since the spurious antisense reads are misassigned sense reads, RoSA then adds the spurious antisense count for each gene to its sense count.

#### 2.4.1 Counting antisense reads

In order to apply the antisense correction factor, counts of antisense reads for each gene are required. Counting antisense reads is not directly supported by read counting tools. However, it can be performed with featureCounts [34] by setting parameters to indicate that reads are stranded in the opposite direction to which they are. Unfortunately, if there are overlapping genes then reads in the overlaps will be counted twice using this tactic. As reads in regions of gene overlap are necessarily ambiguous, they cannot be considered to be antisense, spurious or otherwise. RoSA avoids this issue by building a custom antisense annotation based on the input sense annotation but excluding regions where genes on opposite strands overlap. Different gene transcripts are accounted for by merging all transcripts for a gene into a single *maximal transcript*. Whenever exons of different transcripts overlap in the annotation, the exon in the maximal transcript is the maximum extent of both exons. Given a maximal transcript, the script creates an antisense feature on the opposite strand which runs for the full extent of the maximal transcript. If the maximal transcript of another gene overlaps with the antisense feature, then the antisense feature is truncated to avoid overlapping. Once an antisense annotation is available, a read counting tool can be used to count antisense reads, by providing the antisense annotation instead of the standard annotation.

### 2.5 *Arobidopsis thaliana* datasets with spurious antisense

A procedure to experimentally generate RNA-Seq data with specific levels of spurious antisense is not known. Our main experimental data (Experiment 1) is therefore drawn from the study which originally motivated our investigation into incorrectly assigned antisense reads. In this study, spurious antisense occurred by chance at varying orders of magnitude across different replicates. Additionally, we perform a meta-analysis using three other *Arabidopsis thaliana* datasets (Experiments 2-4) and data from ENCODE (detailed in the Supplementary Material).

#### 2.5.1 Sample preparation and sequencing

The RNA-Seq data for Experiments 1-4 are wild-type (WT) *Arabidopsis thaliana* Colombia-0 (Col-0) biological replicates. WT *A. thaliana* Col-0 seeds were sown aseptically on MS10 plates. The seeds were stratified for 2 days at 4°C and then grown at a constant 21°C under a 16-h light/8-h dark cycle for a further 14 days, at the end of which the seedlings were harvested. Total RNA was isolated from the seedlings with the RNeasy Plant Mini Kit (Qiagen). In Experiment 1, DNAse digestion was performed on column, as a part of RNA isolation, and 8 μl of ERCC spike-ins (External RNA Controls Consortium 2005) at a 1:100 dilution was added to 4 μg of total RNA. Libraries were prepared according to the TruSeq® Stranded mRNA Sample Preparation Guide Rev E. In Experiments 2-4 the samples were treated with TURBO™ DNase (Ambion) and 4μ1 of ERCC spike-ins at a 1:100 dilution were added to 1μg/6μl of total RNA. Libraries were prepared using the Illumina TruSeq Stranded Total RNA with Ribo-Zero Plant kit. In all experiments, the libraries were sequenced on a HiSeq2000 at the Genomic Sequencing Unit of the University of Dundee. Experiment 1 has 3 replicates, processed as one batch, with a total of 4 × 10^8^ 150-bp paired-end reads. Experiments 2 and 3 have 7 biological WT replicates, while Experiment 4 has 3, for a total of 17 biological WT replicates and ~1.7 × 10^9^ 100-bp paired-end reads across the three experiments. The same lab sowed, grew and harvested the plants, and prepared the libraries. The sequencing was performed on the same machine by the same people at the same sequencing facility and all the samples include the ERCC spike-ins which can verify the WT samples are consistent and comparable across experiments.

#### 2.5.2 Quality Control, Alignment and Quantification

The quality of the data was quantified using FastQC v0. 11.2 [36] with all the replicates performing as expected for high quality RNA-Seq data with good median per-base quality (≥28) across >90% of the read length. The read data for all experiments were aligned to the TAIR10 [37] *Arabidopsis thaliana* genome using the splice-aware aligner STAR v2.4.2a [38] for Experiment 1 and STAR v2.5.0 for Experiments 2-4. The index was built with `–sjdbOverhang 149` (Experiment 1) or `–sjdbOverhang 99` (Experiments 2–4) and the alignment was run with parameters: `–outSJfilterlntronMaxVsReadN 5000 10000 15000–outSAMAttributes All–outFilterMultimapNmax 2–outFilterMismatchNmax 5–outFilterType BySJout`.

The read data were also aligned to the ERCC spike-ins annotation, using the same parameters. Read counts per gene were then quantified from these alignments with featureCounts v1.5.0-p1 using the publicly available TAIR10 annotation with the parameters: -s 2 -p -t exon– `largestOverlap`. After running RoSA’s *make_annotation* script to build an antisense annotation, antisense read counts per gene were quantified in the same way, with the parameters: `-s 2 -p -t antisense–largestoverlap`. Finally, spliced sense and antisense reads were counted using RoSA’s *count_spliced* script with the TAIR10 annotation.

## 3 Results

We used RoSA to analyse our data from Experiment 1 for spurious antisense, using both the spike-in and spliced reads counts. RoSA calculated antisense:sense ratios for the spike-ins (**Figure *2***), showing that the 3 replicates have antisense:sense ratios on the spike-ins of 0.0008, 0.004 and 0.011. Although these ratios are small, if the replicates were being compared for differential expression, the differences are potentially substantial for highly expressed genes, and could lead to differential antisense expression being called erroneously.

**Figure 2:**
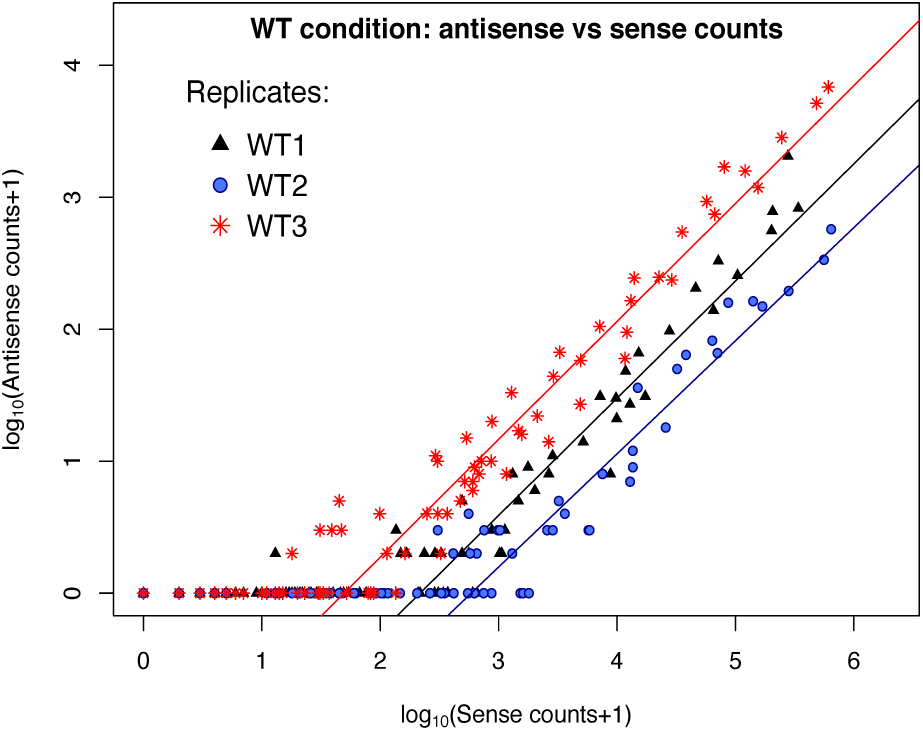
Antisense versus sense counts for the ERCC spike-ins for each replicate in Experiment 1. Points represent antisense and sense read counts for individual spike-ins. Each line is the average antisense: sense ratio for one replicate. Here, antisense:sense ratios vary by an order of magnitude across the 3 replicates, with values of 0.004 (WT1), 0.0008 (WT2), and 0.011 (WT3).

For each replicate we calculated the spurious antisense:sense ratios for the spliced reads with RoSA, and compared them to the spike-ins. An overview of the results for all three replicates is shown in **Figure *3*. Error! Reference source not found**. (Row 1) shows the plots of spliced read counts and spike-in counts per gene or spike-in, for each replicate, demonstrating that the spurious antisense levels calculated from the spike-ins are in good agreement with the levels calculated from the spliced reads.

**Figure 3:**
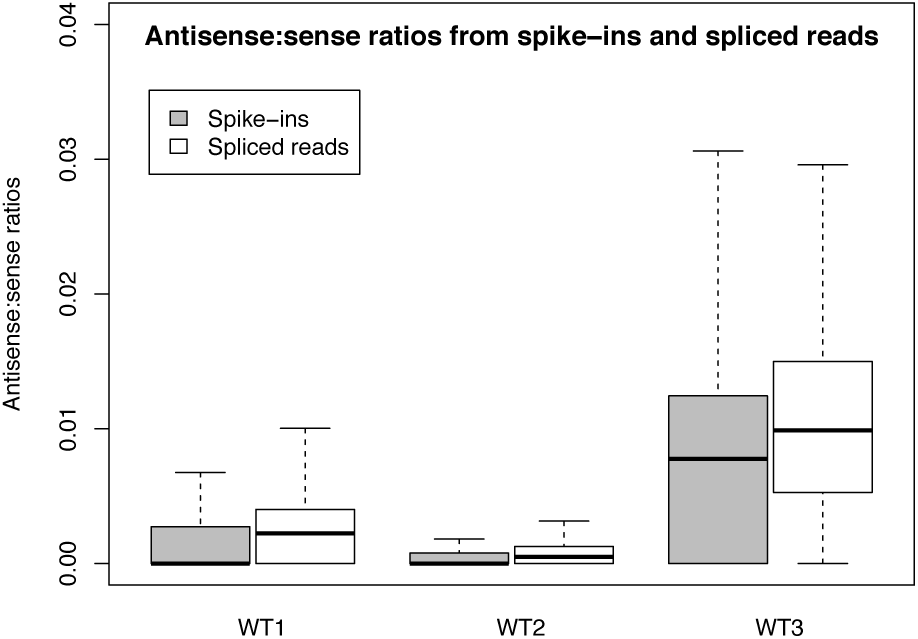
Comparison of antisense:sense ratios calculated from spliced reads or spike-ins, by replicate. Ratios estimated from spike-ins show good agreement with ratios estimated from spliced reads. Outliers not shown.

Finally, RoSA calculated a spurious antisense correction across the whole of each replicate. Every spliced gene was corrected with the antisense:sense ratio specific to the gene, and unspliced genes were corrected using the mean ratio calculated from the spike-ins. (RoSA also allows the unspliced correction to be calculated from the mean spliced reads ratio, for datasets without ERCC spike-ins). Overall, RoSA reduces the correlation between antisense and sense counts in the data (**Error! Reference source not found**., Rows 2 and 3), as would be expected with a reduction in incorrectly assigned reads. Two examples of corrections made by RoSA are shown in **Figure *4***: where the antisense signal appears to be almost entirely spurious, RoSA’s correction factor reduces the antisense counts substantially, but where there also appears to be some real antisense signal, RoSA’s correction factor leaves a higher proportion of counts.

**Figure 4:**
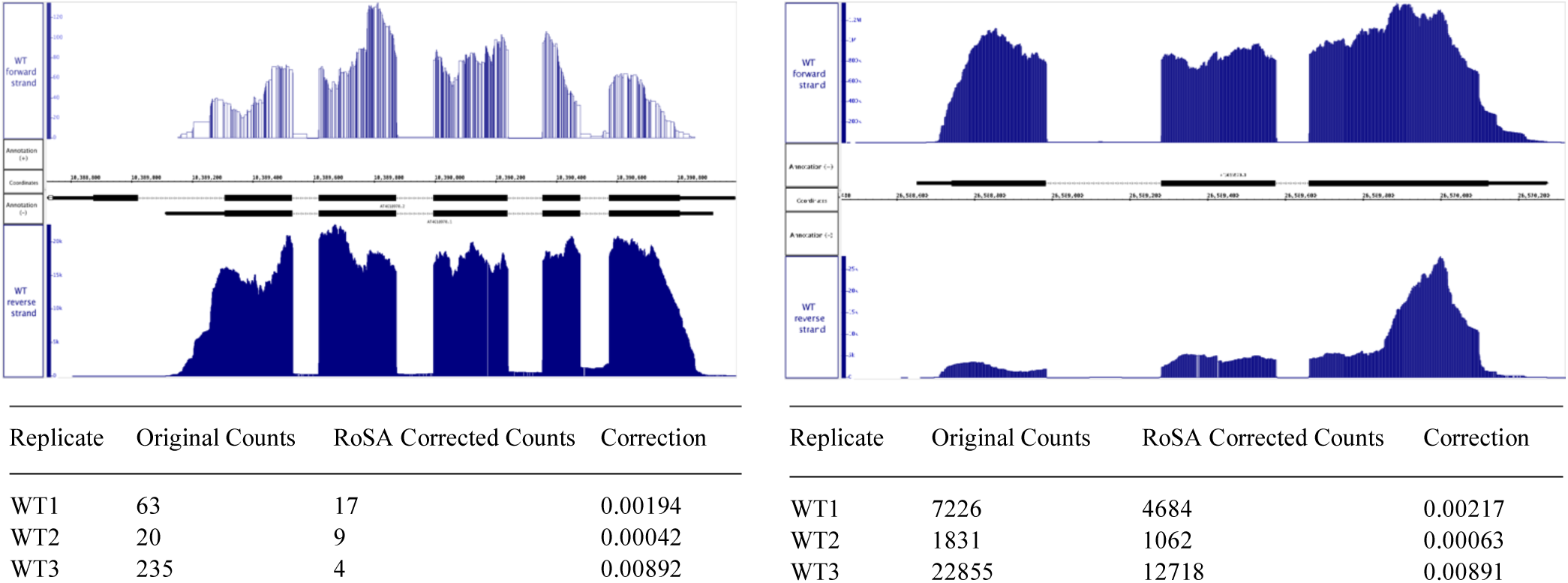
Two genes with differing antisense expression profiles and the read count corrections proposed by RoSA. The reverse strand gene AT4G18970 (left) has antisense expression which clearly matches the splice sites of the sense strand. RoSA eliminates almost all of the antisense reads. The forward strand gene AT5G66570 (right) has both antisense expression matching the sense strand splice sites, and a peak at the 5’ end which is unlikely to have resulted from incorrect read assignment. RoSA only reduces the antisense counts by around 40%. (Figures generated by IGB [1])

As well as identifying instances of antisense expression, looking at antisense counts in this way may also be useful in identifying misannotated genes. For example, in our data there are many genes where the antisense:sense ratio is more than 1 (e.g. see points lying above *x=y* in **Error! Reference source not found**., Row 2), which may be due to the wrong strand being assigned to genes in the annotation used.

### 3.1 Comparing antisense: sense ratios

Calculating antisense:sense ratios allows comparisons of spurious antisense to be made between replicates and between experimental conditions, and can reveal whether there are systematic differences which might confound experimental comparisons. For example, **Figure 1** presents results from an RNA-Seq experiment where spurious antisense levels differed by an order of magnitude between replicates. In this experiment, the WT replicates had spurious antisense:sense ratios of 0.0031 (SD 0.0116), 0.0009 (SD 0.0070) and 0.0111 (SD 0.031).

To determine the extent of this problem for RNA-Seq datasets in general, we investigated the spurious antisense levels across a range of experiments and research groups. We analysed antisense reads assigned to the spike-ins from three other experiments in our lab (Experiments 2–4), as well as 195 publically available human datasets from the ENCODE project (see Supplementary Material) that included the ERCC spike-ins [29]. A separate antisense:sense ratio was calculated for each replicate in each experiment, (**Figure *5***) showing that spurious antisense reads are present at varying levels and can range across several orders of magnitude. This presents a serious quality control issue for anyone investigating differential antisense expression: a real difference in antisense expression could be completely masked by a difference in spurious antisense.

**Figure 5:**
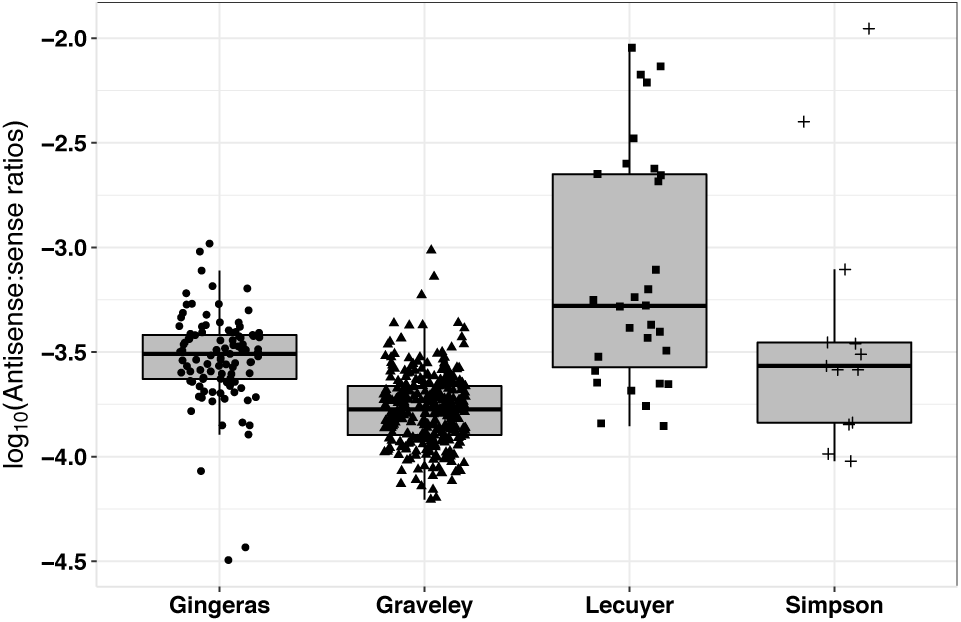
Spurious antisense:sense ratios for spike-ins, by research group. Data are from either from ENCODE (Gingeras, Graveley and Lecuyer) or our own group (Simpson). Each point represents the ratio for a replicate. Ratios range from 0.0111 to 0.00003

**Figure 6:**
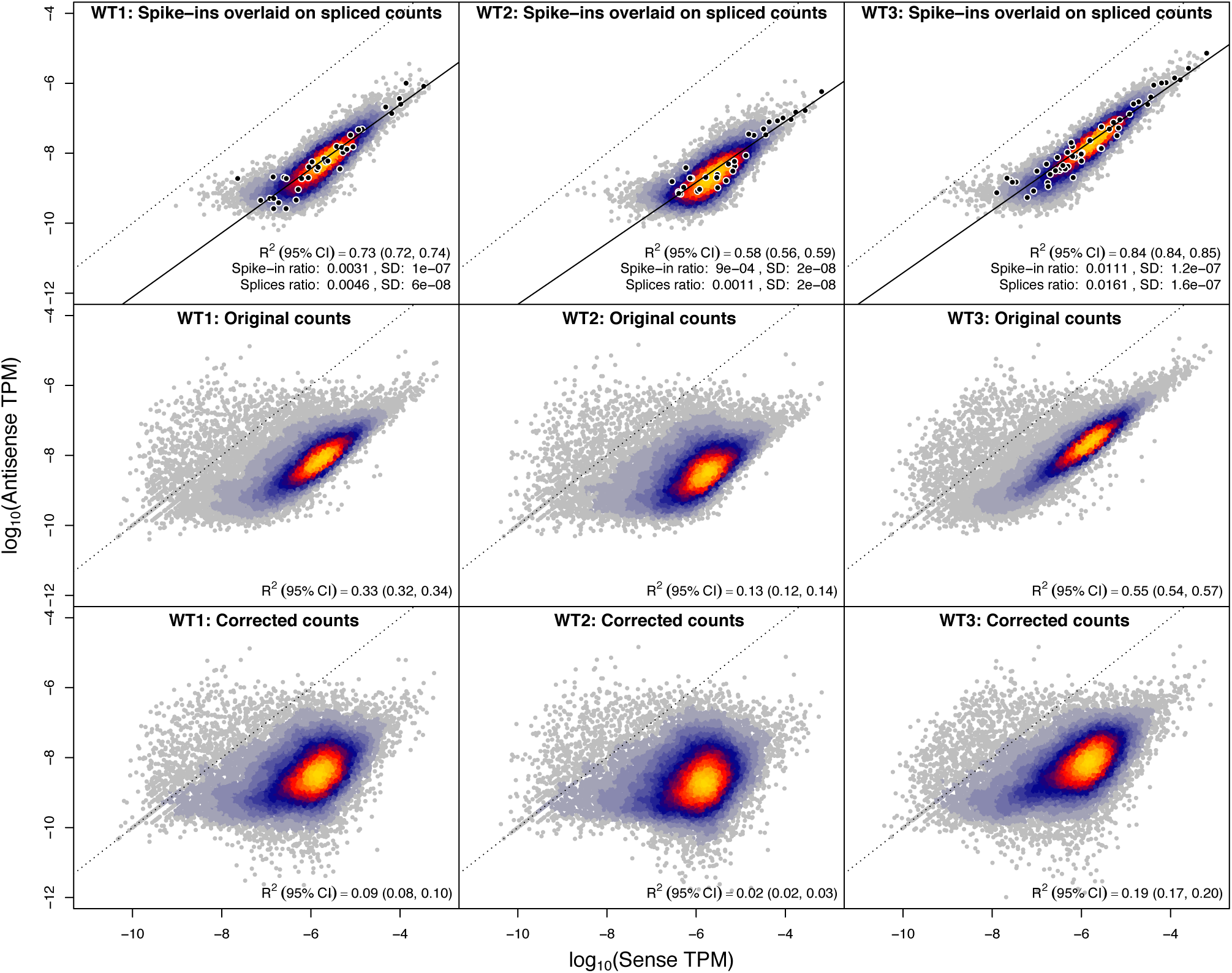
Normalised antisense versus sense counts by replicate. **Each** column presents data for one replicate. Row 1: Comparison of antisense:sense ratios calculated from spike-ins and from spliced reads. Black points and fit line correspond to spike-in values; the heat map plots normalised antisense and sense counts for each spliced gene. The antisense:sense ratios calculated from the spike-ins or spliced reads are in good agreement. Row 2: The original antisense counts are correlated with the sense counts (plotted for all genes). Row 3: The corrected antisense counts show much weaker correlation. On all plots the dashed line marks y=x; points above this line correspond to genes where the antisense:sense ratio is > 1.

## 4 Conclusions

Spurious antisense is common in strand-specific RNA-Seq datasets, and can occur at varying levels across replicates in the same experiment. Differing levels of such incorrectly assigned reads are enough to disrupt differential expression analyses of antisense gene expression.

We have developed a new tool, RoSA, which can detect, quantify and correct for spurious antisense. RoSA provides an important quality control step for researchers analyzing antisense expression in their data.

